## Supplementary Information for "Plasma exosomes from individuals with type 2 diabetes drive breast cancer aggression in patient-derived organoids"

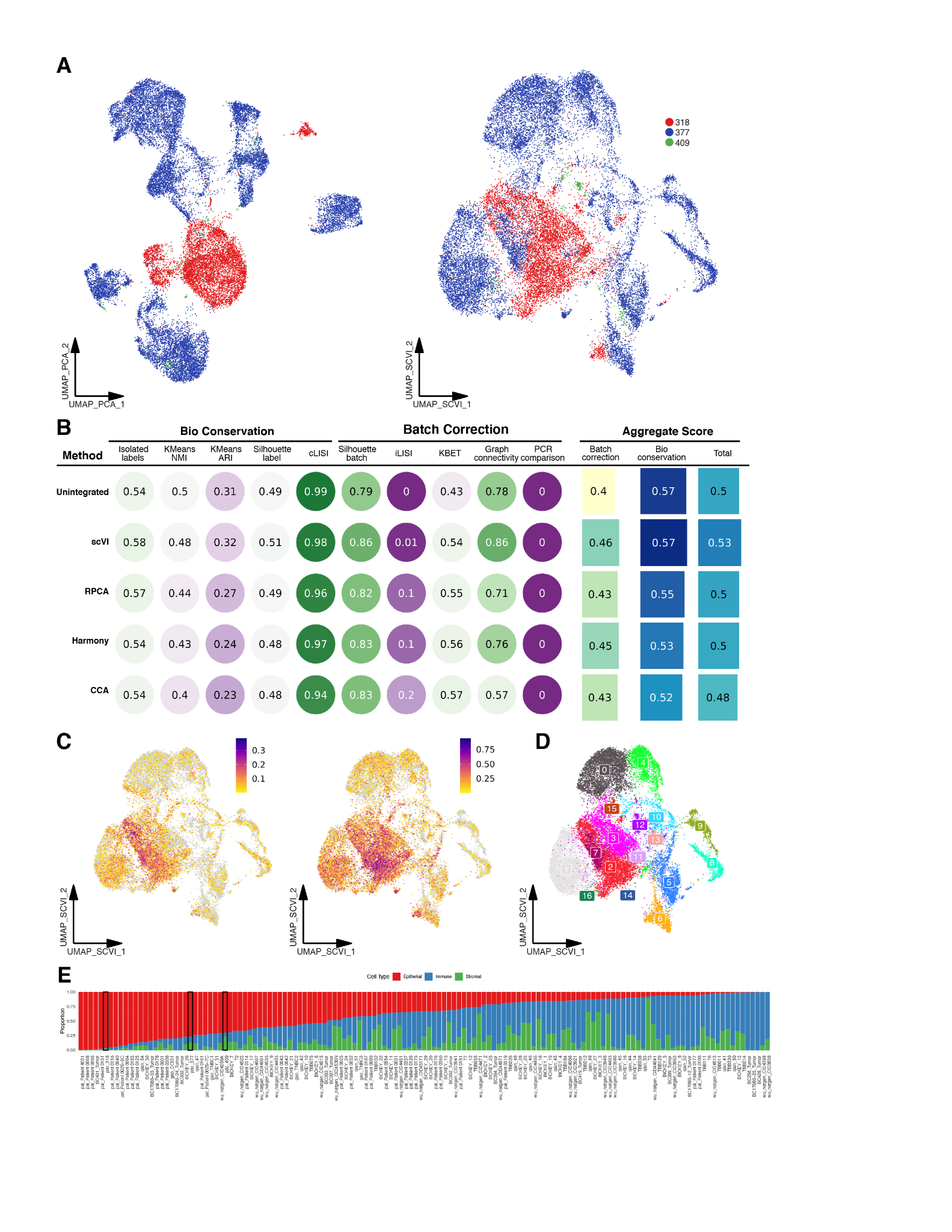

**Fig S1. scRNAseq metrics and non-integrated data.**

**A.** UMAP visualizations of PDOs colored by patient, showing integrated (left) and scVI-integrated (right) data. **B.** Benchmarking results assessing batch correction performance across dataset. **C.** UMAP visualizations colored by module score of COSMIC CGC (left) and GOBP_MAMMARY_EPITHELIAL_PROLIFATION (right) gene expression. **D**. UMAP visualization of scVI-integrated data colored by Louvain clusters. **E.** Barplot showing proportion of cells identified as epithelial, stromal, or immune in benchmarking atlas. Distributions of PDOs boxed.

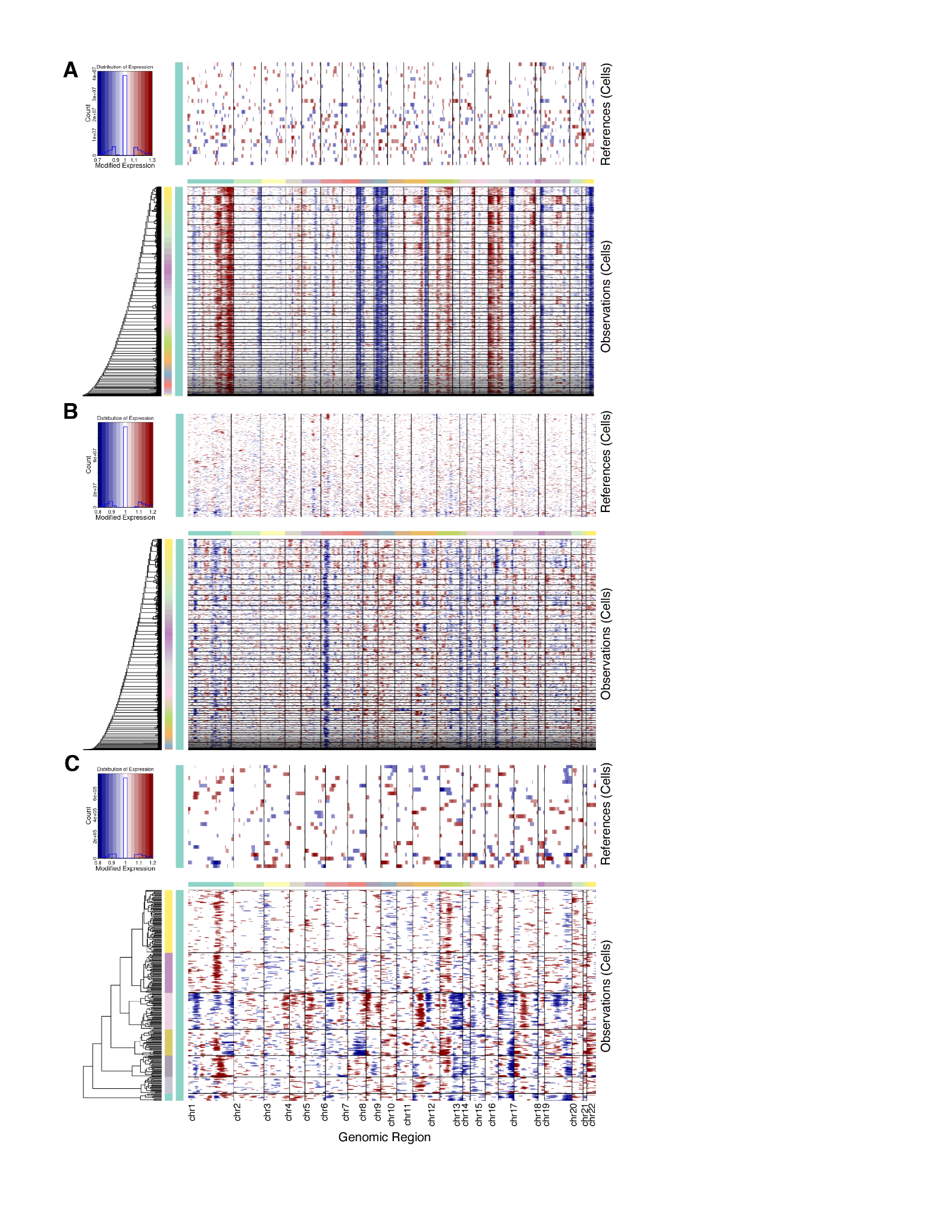

**Fig S2. Identification of malignant cells.**

**A/B/C.** InferCNV heatmaps of all malignant cells for patients 318 (**A**), 377 (**B**) and 409 (**C**). Immune cells used as reference on an individual patient basis.

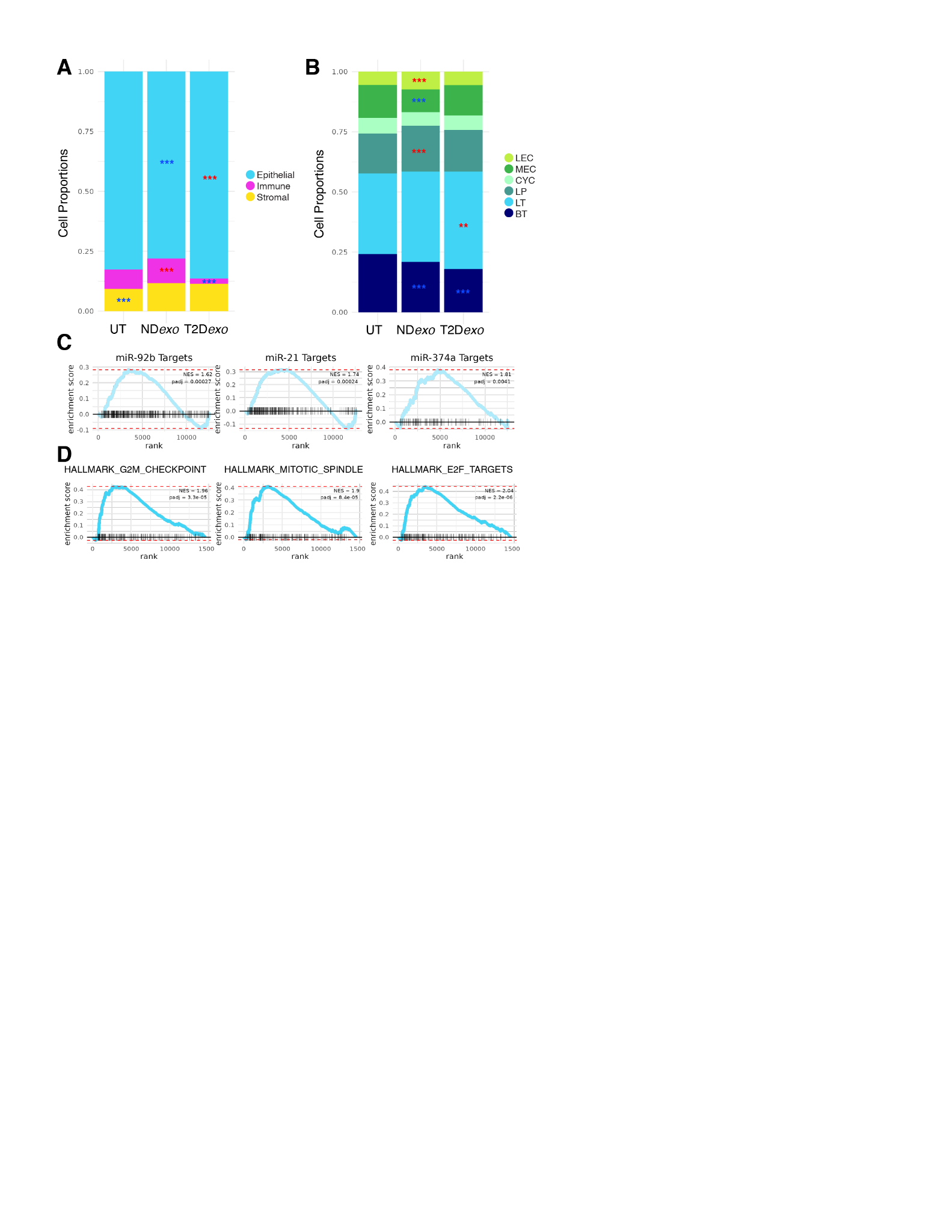

**Fig S3. Relative abundance and cell type dynamics.**

**A.** Relative proportion of coarse annotations per treatment. Note uniformity of stromal compartment across treatments. Significance calculated via binomial linear regression model, ***<0.001, **<0.01, *<0.05. **B.** Relative proportion of all epithelial and tumor subclones. Significance calculated via binomial linear regression model, ***<0.001, **<0.01, *<0.05. **C/D.** GSEA enrichment plots for representative gene sets upregulated in LT1 (**C**) or LT3 (**D**).

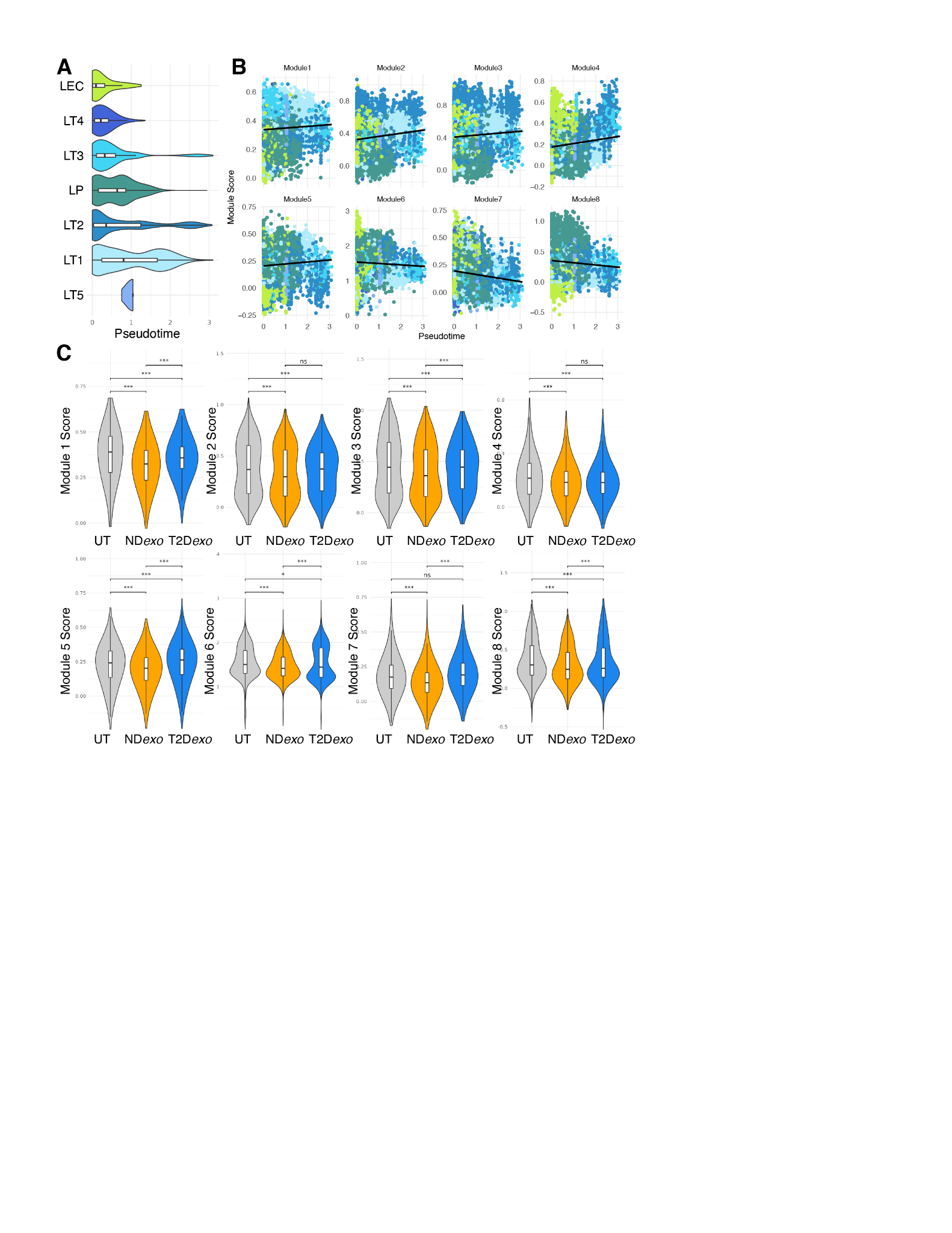

**Fig S4. Pseudotime analysis and module characterization.**

**A.** Violin plots of pseudotime calculated for luminal-like epithelial cells per cell type. **B.** Scatter plots of module gene set expression as a function of pseudotime. Dot color denotes epithelial cell subtype assigned to cells. Black line showing linear regression of best fit. **C.** Violin plots of module expression in luminal-like epithelial cells per treatment. Significance calculated via linear mixed effects model, ***<0.001, **<0.01, *<0.05.

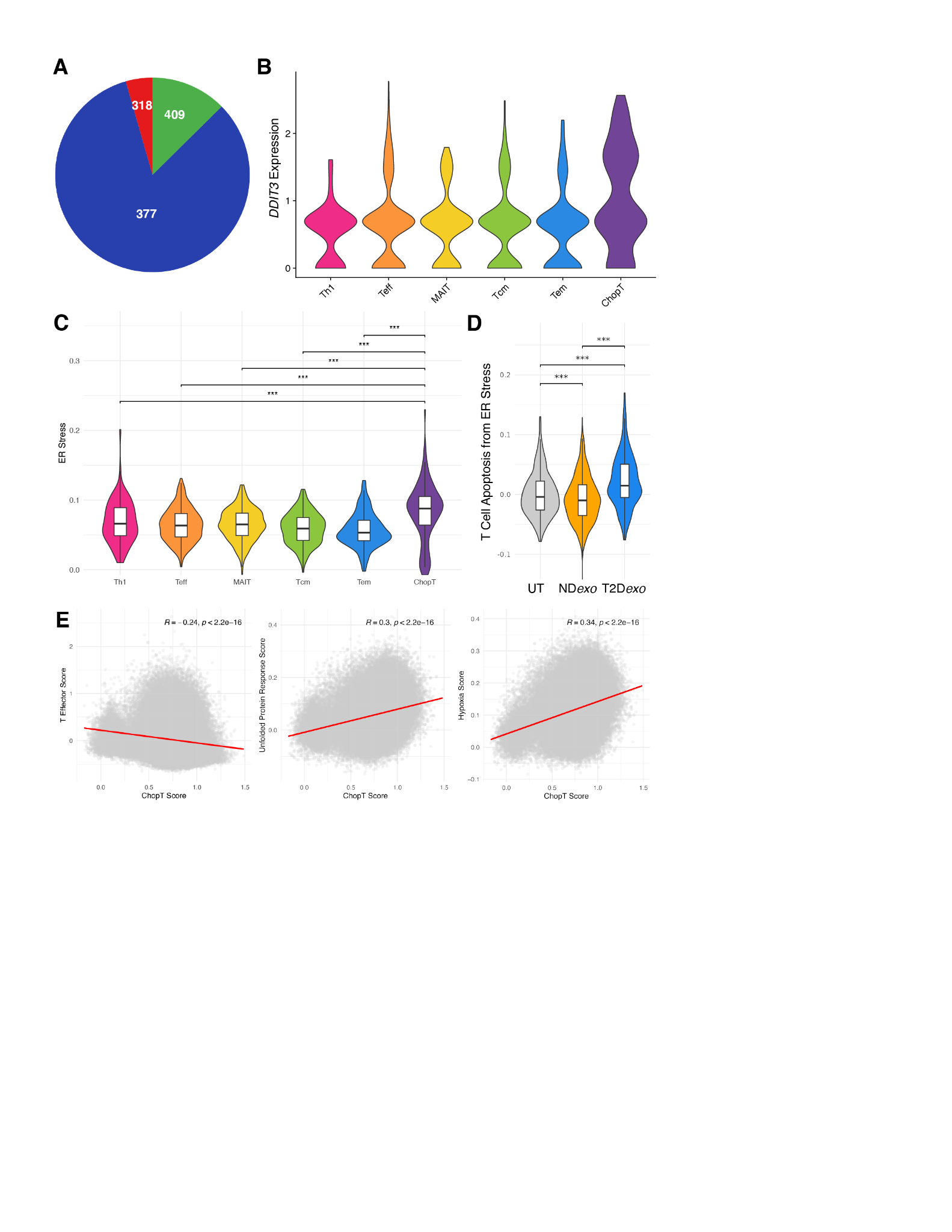

**Fig S5. Characterization of ChopT cells.**

**A.** Pie chart of the distribution of ChopT cells across patients, demonstrating its conservation. **B.** Violin plots of *DDIT3* expression across T cell states. **C.** Violin plots of expression of GOBP_RESPONSE_TO_ENDOPLASMIC_RETICULUM_STRESS across T cell states. Significance calculated via linear mixed effects model, ***<0.001, **<0.01, *<0.05. **D.** Violin plots of expression of GOBP_INTRINSIC_APOPTOTIC_SIGNALING_PATHWAY_IN_RESPONSE_TO_ENDOPLASMIC_RETICULUM_STRESS in T cells across treatment groups. Significance calculated via linear mixed effects model, ***<0.001, **<0.01, *<0.05. **E.** Scatter plots of ChopT module score versus effector score (left), UPR score (middle), and hypoxia (right) across the pan-breast cancer immune cell atlas. Red lines indicate lines of best fit. Spearman correlation coefficients and p-values shown.

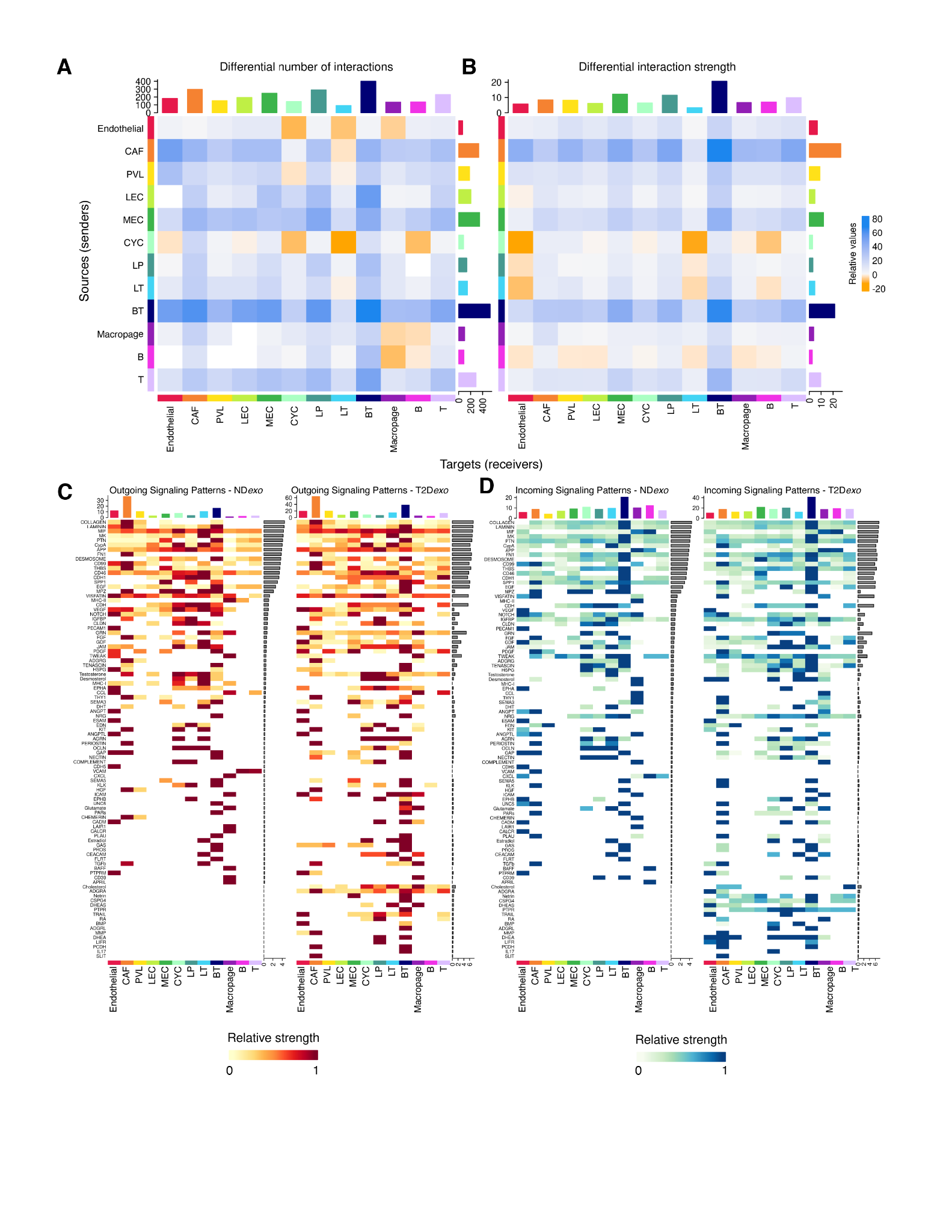

**Fig S6. Cell-cell communication networks.**

**A**. Heatmap showing the number of interactions among different cell types. The blue (orange) colored edges represent increased (decreased) signaling in T2D*exo* compared to ND*exo*-PDOs. **B**. Heatmap showing the differential interaction strength among different cell types. The blue (orange) colored edges represent increased (decreased) interaction strength in T2D*exo* compared to ND*exo*-PDOs. **C.** Comparison heatmap of patterns of ingoing communication in ND*exo*-PDOs (left) and T2D*exo*-PDOs (right). **D.** Comparison heatmap of patterns of outgoing communication in ND*exo*-PDOs (left) and T2D*exo*-PDOs (right).

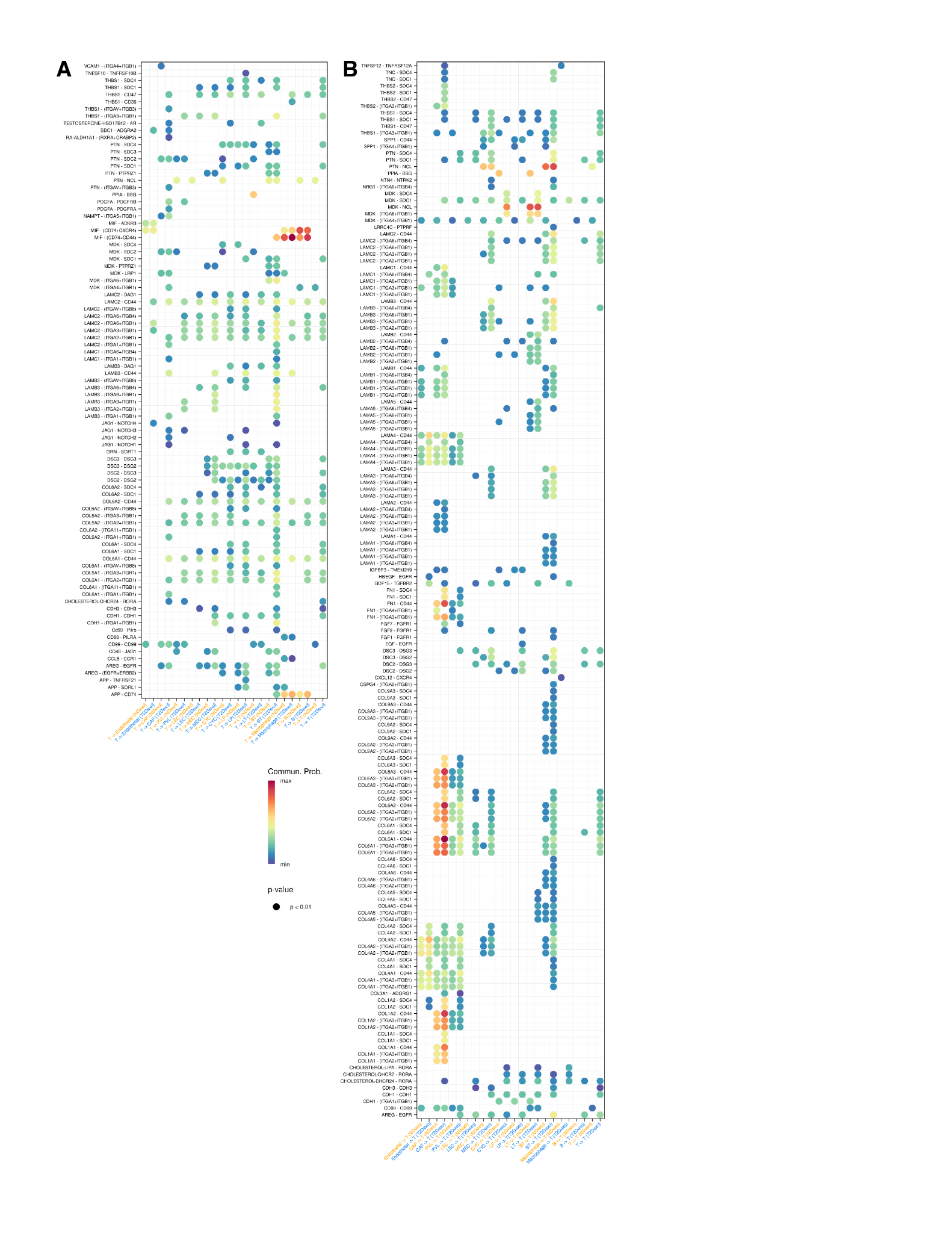

**Fig S7. T cell-specific communication networks.**

**A/B.** Dot plot of expression levels of significant (p < 0.01) ligand-receptor pairs from all senders to (**A**) and all receivers from (**B**) T cells. *P* values are computed from a one-sided permutation test according to CellChat.

**Table S1. Clinicopathological characteristics of patient tumors.**

This table summarizes the age at diagnosis, tumor size, histological type, tumor grade, lymph node involvement, TNM stage, hormone receptor status (ER, PR), HER2 status, and Ki-67 proliferation index for each patient included in the study.

**Table S2. Marker genes for broad cell types identified in PDOs.**

This table lists the marker genes used to identify and characterize broad cell types maintained in PDOs. The table includes gene names per cell type and associated statistics.

**Table S3. GSEA results comparing T2D*exo* vs. ND*exo* across all cells.**

This table lists the significant gene sets (adjusted p-value < 0.05) identified by GSEA when comparing all cells captured within T2D*exo*-PDOs vs. ND*exo*-PDOs. The table includes the gene set names and associated statistics.

**Table S4. Differentially expressed genes in T2D*exo* vs. ND*exo* across all cells.**

This table lists the differentially expressed genes identified in T2D*exo*-PDOs compared to ND*exo*-PDOs across all cell types with associated statistics.

**Table S5. Circularity measurements of PDOs.**

This table summarizes the area, perimeter, and circularity measurements of individual PDOs. Singlets were removed during thresholding.

**Table S6. Composite gene signature for survival analysis.**

This table lists top genes upregulated in either T2D*exo*-treated PDOs or in ND*exo*-treated PDOs, which together comprise the composite signature applied to TCGA and METABRIC cohorts.

**Table S7. Marker genes for epithelial clusters identified in PDOs.**

This table lists the marker genes used to identify and characterize epithelial and tumor subclones maintained in PDOs. The table includes gene names per cell type and associated statistics.

**Table S8. GSEA results comparing LT1 vs. all other luminal epithelial cells.**

This table lists the significant gene sets (adjusted p-value < 0.05) identified by GSEA when comparing LT1 cells compared to all other luminal-like epithelial cells. The table includes the gene set names and associated statistics.

**Table S9. Differentially expressed genes along pseudotime.**

This table presents the results of graph autocorrelation analysis, identifying genes with significant expression changes along the pseudotime trajectory. The table includes gene names and associated statistics.

**Table S10. Genes comprising co-regulated modules in pseudotime.**

This table lists the genes grouped into distinct modules of co-regulation. Each module represents a cluster of genes with synchronized expression patterns along the pseudotime trajectory, reflecting their potential involvement in specific biological processes during tumor progression.

**Table S11. Marker genes for immune clusters identified in PDOs.**

This table lists the marker genes used to identify and characterize immune and T cell states maintained in PDOs. The table includes gene names per cell type and associated statistics.

**Table S12. Differentially expressed genes in T2D*exo* vs. ND*exo* across immune cells.**

This table lists the differentially expressed genes identified in T2D*exo*-PDOs compared to ND*exo*-PDOs across immune cell types with associated statistics.

**Table S13. GSEA results comparing T2D*exo* vs. ND*exo* across immune cells.**

This table lists the significant gene sets (adjusted p-value < 0.05) identified by GSEA when comparing immune cells captured within T2D*exo*-PDOs vs. ND*exo*-PDOs. The table includes the gene set names and associated statistics.

**Table S14. Enrichment results on branch point via K2Taxonomer.**

This table summarizes the significantly upregulated gene sets (p-value < 0.05) identified during K2Taxonomer analysis of ChopT vs. normal development branch point. The table includes the gene set names and associated statistics.

**Table S15. Intercellular communication inferred from ligand-receptor interactions.**

This table provides an overview of the cell-cell communication networks within T2D*exo*-PDOs and ND*exo*-PDOs as inferred from ligand-receptor interactions. This table lists identified ligand-receptor pairs, their pathway annotations, the cell types involved in these interactions, and associated statistics.
